# Intriguing relationship between chromosome I nuclear aneuploidy and mitochondrial intron deletion in yeast

**DOI:** 10.1101/2025.05.25.656015

**Authors:** Ali Gargouri

**Affiliations:** Centre de Génétique Moléculaire, CNRS Gif sur Yvette, France

## Abstract

A curious phenomenon was discovered when examining revertant colonies of the M3041 mutant located in the second intron of mitochondrial cytochrome b gene of *Sacchromyces cerevisiae*. Several colonies carrying a sector appeared on selective glycerol medium and all correspond to a reversion of M3041 by intron deletion. Colonies of revertants resulting from classical reversion were never sectored. The cells of the sectors, called S, presented a very typical growth profile on glycerol (selective medium for respiration) which differs from that of the rest of the colony called AS for anti-sector. The profiles are identical on glucose-based medium. Molecular analysis, by Pulsed Field Gel Electophoresis, of the S cells showed that they had lost one copy of chromosome 1, suggesting that the original a-haploid mutant had diploidized into mat (a/a). The aneuploid state was validated by crossing with a genetically marked strain followed by mass spore analysis. We then asked whether the aneuploid genetic background would influence the frequency of reversion by intron deletion. This was achieved through the conversion of S and AS cells to rho^0^ followed by cytoduction of mitochondria carrying M3041 or W91 mutations, knowing that W91 never reverses by intron deletion. Interestingly, there are significantly more revertants (by intron deletion) in the S than AS context for M3041, whereas W91 reversed with the same frequency in both contexts. Nuclear aneuploidy therefore promotes mitochondrial reversion by intron-deletion. We conclude that there is a particular relationship between nuclear aneuploidy and intron deletion during the reversion of mitochondrial intron mutations in yeast.

## Introduction

The yeast *Saccharomyces cerevisiae* is an aerobic-facultative organism that respires through mitochondria and ferments glucose into ethanol (*1*). Yeast mitochondria include circular mitochondrial DNA which is larger than that of mammals because of large inter-genic spaces and especially the introns (*2*,*3*). In addition, these introns have several characteristics that distinguish them from the nuclear introns of yeast or higher organisms. All nuclear introns begin with the dinucleotide GT and end with AG (*4*), never code, have no determined secondary structure and are generally excised in the same way thanks to the spliceosome (*5*). On the other hand, the mitochondrial introns do not have the same di-nucleotides at the ends, have two different secondary structures group I and group II, some are able to self-excise from the pre-messenger RNA (*6*,*7*). Others require the action of protein cofactors and some of them code for a certain number of proteins. They can in fact code for a protein called maturase involved in the splicing of the intron that encoded it (*8*,*9*). They also encode other proteins such as reverse transcriptase (*10–12*), involved in the intron mobility and the recombinase/endonuclease involved in the homing phenomenon (*13*,*14*). Along with mitochondrial introns, a particular intron in the Tetrahymena nucleus has generated much curiosity since it was found to be capable of self-excise(*15*,*16*).This intron also had the group II structure. This self-excision property was associated with yeast mitochondrial introns even if some continue to require the intervention of maturase in their excision *in vivo*. Subsequently, these structural (groups I and II), self-sxcision and coding properties of maturase were found in other kingdoms such as bacteria and plants(*17*). There are even maturases which are encoded in the nucleus of certain plants and which will then be sent to the mtochondria to excise introns (*18*).

The first respiratory mutants were discovered in yeast with the famous "petite" mutants that are irreversible because they consist of large deletions of mitochondrial DNA (*19*). Later, reversible mitochondrial mutants were discovered, called mit-, and correspond to single mutations (missense, nonsense, or frameshift) taking place in the genes coding for a subunit of the respiratory chain, in particular, the cytochrome b gene (cyt.b) and the genes encoding the three cytochrome oxidase subunits (cox) (*19–21*). With the discovery of introns, mit- mutations encompassed exon mutations and intron mutations. The latter could take place either in the coding phase of the intron, which is often fused in phase to the exon that preceded it, or in the non-coding phase of the intron which generally disrupts the secondary structure of the intron in question. As simple mutations, they can be reversed by different modes of classical reversion; One can cite the change of the mutated base into another base restoring either the original sequence (true reversion) or a similar and functional sequence (pseudo reversion) (*22*). There are also cases of reversion where the mutation is maintained but is suppressed by informational or functional suppressors (*22–24*).

Informational suppressors affect the factors involved in ribosomal translation (rRNA, ribosomal protein, etc.) which will therefore allow correct reading of the sequence. These suppressors are specific for mit- having occurred in the coding part of the intron. We can cite the mim3-1suppressor (*23*) which modified the 16S rRNA (*24*) and nam3-1 which affected the Mitochondrial Release Factor (*25*). Functional suppressors can act on mit- taking place in the coding or non-coding phase. We can cite the MRS3 suppressor (*26*), also called NAM10 (*27*), which acts on both types of mutations (*27*) and corresponds to an iron transporter in the mitochondrial membrane (*26*).

Another mode of reversion has been discovered for mutations in yeast mitochondrial introns and in particular in the cytochrome b gene: this is the deletion of the mutated intron which is most often associated with deletion of other introns (*27*). Indeed and by as illustrative example, in the case of a mutation in the second intron of cyt.b, called bi2, the majority of deletion revertants have lost either the bi1 and bi2 introns or the bi1, bi2 and bi3 introns (*28*,*29*). This phenomenon of reversion by intron deletion was shown to involve a reverse transcription step because it cannot take place in the absence of the first two introns ai1 and ai2 of the cox1 gene (subunit 1 of cytochrome oxidase) (*12*,*29*). Note that these two introns are homologous to viral RT (*10*) and were shown to possess reverse transcriptase activity (*11*). The presence of both introns or one of them is sufficient to allow reversion by intron deletion (*12*, *29*).

This phenomenon of reversion of mit- mutations by intron deletion (ID) has been particularly studied in bi2 intronic mutations. These are mutations in the coding or non-coding part of the bi2 intron, recalling that structurally this intron belongs to group I introns. Interestingly, some mutations can reverse through this phenomenon as well as by classical reversion and extragenic deletion, the most representative being the M3041 mutation (*27–29*). Other mutations cannot be reversed by intron deletion, such as the W91 mutation, although it is very close to M3041 (*27–29*).

Special mention should be given to the M3041 mutation in the bi2 intron which is a frame- shift +A mutation, creating a stop codon immediately where the A is inserted. Indeed, it can be suppressed by informational suppressors such as mim3-1 in the mitochondrial 16S rRNAgene (*27*), by nam3-1 (*23, 27*) and finally by the functional suppressor MRS3/NAM10 (*26,27*). More interestingly, it reverts very efficiently by deletion of the introns.It has therefore always been taken as a model for the study of each mode of reversion and in particular the deletion of introns (*27–29*). The majority of deletions concerned either the introns (bi1 + bi2) or the (bi1 + bi2 + bi3) introns (*28, 29*), recalling that the intron bi4 is never deleted as it is required for the splicing of the ai4 intron in the cox1 pre-mRNA (*30*). This work describes a rare phenomenon discovered during the study of M3041 revertants. It began with the observation of sectors on revertant colonies and led us to discover a curious relationship between the deletion of introns in the mitochondria and an aneuploidy of chromosome 1 in the yeast nucleus.

## Results

### Discovery of the sectored colonies

This phenomenon was discovered on several revertants of the haploid strain CK247 (M304l) and appeared on Glycerol (N3) plates after prolonged incubation at 28°C. These are sectored colonies always having the same type of sector, Fig.1, with the following characteristics:

**Fig. 1:**
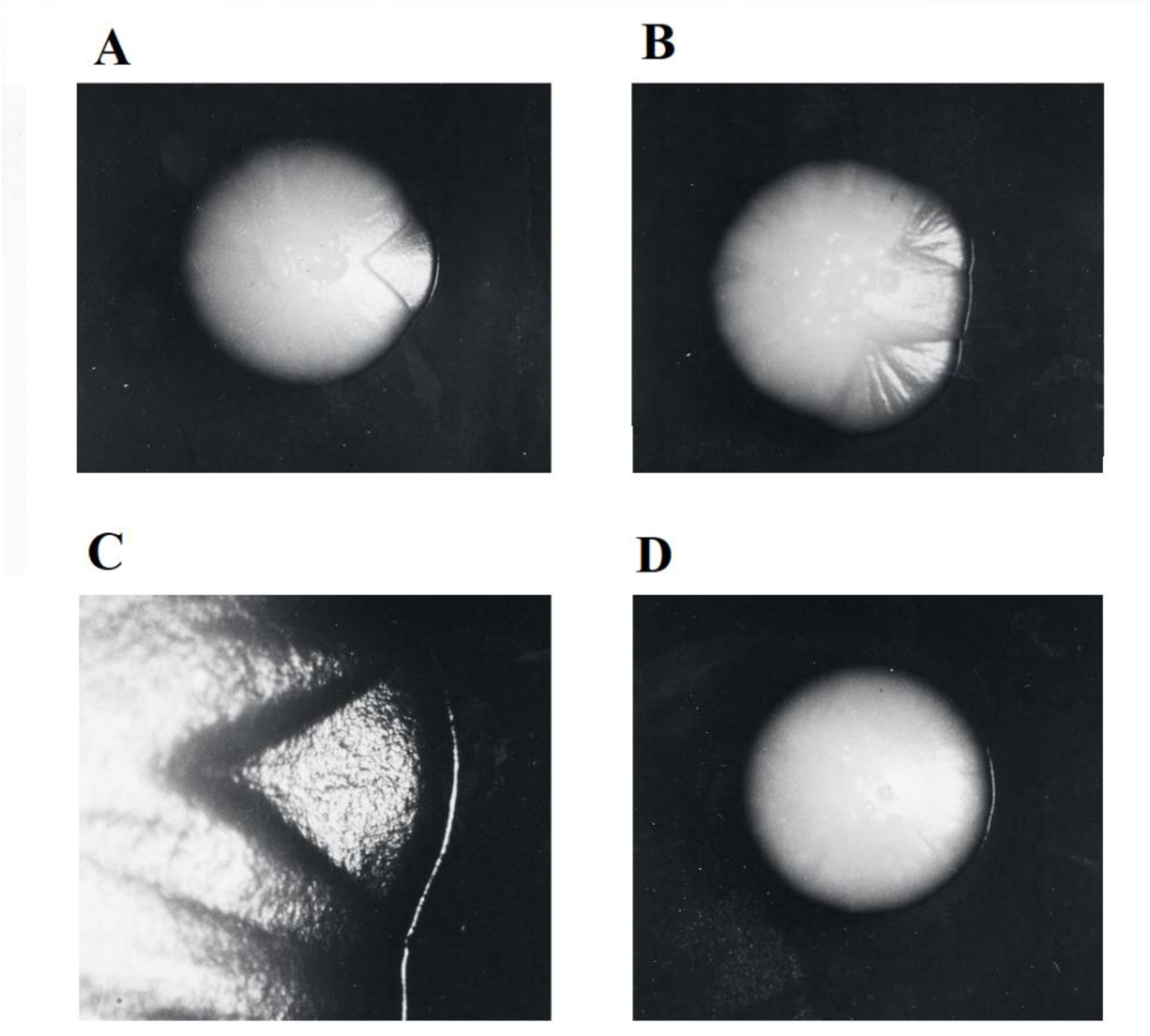
Phenotype of sectored and non-sectored colonies. After 15 days of incubation at 28°C of the CK247 strain (carrying the M3041 mutation) spread on Glycerol N3 medium, revertant colonies appear and some of them were sectored. The photos are taken with the same magnification with the exception of the photograph denoted C. With: **A** sectored colony with one sector; **B** sectored colony with two sectors; **C** detail of a sector region; **D** non-sectored colony NS.

- The sector will be called (S) and the rest of the colony anti-sector (AS). The colony with no sector is called non-sectored (or NS).

- The arc formed by the sector is protuberant and sunken compared to the rest of the colony.

- They do not start from the center of the colony except in one case out of nearly 300 colonies observed. This means that it always occurs after the reversion event is triggered.

The average frequency of the *sector event* per cell can be approximately estimated at l0^-5^-10^-6^ (the number of cells accumulated at the time of the emergence of the sector). This frequency is higher than the point mutation frequency and is more reminiscent of a phenomenon of nuclear genetic instability.

During the first observation, five colonies out of thirty analyzed molecularly were sectored (one of which presented two sectors and the others a single sector). The thirty revertants were all of the delta or deletion type: having reversed by loss of introns at the mitochondrial DNA level, two of the colonies in the sector were delta-bil+bi2+bi3 the other three were delta- bil+bi2. We should recall here that it is known that M3041 (located in bi2 intron of the cyt.b gene) reverts by classical reversion and by intron deletion (*28*, *29*). The most frequent events are the deletion of bi2, the mutated intron, but also other introns: bi1+bi2 or bi1+bi2+bi3 and very rarely bi2 alone or bi2+bi3 (*28*, *29*).

### Looking for a possible relationship between *sectorization* and intron deletion?

To answer the question of the relation between sectorization and intron deletion, two other bi2 mutants were used with M3041: W91 (haploid strain CK239) and M2075 (CK312) which are unable to revert by intron deletions (*28*,*29*). Thus, the three strains CK247, CK239 and CK312 were plated out at a density of 5.10^8^ cells/plate on N3 medium and incubated for 15 days at 28°C. Table 1 summarizes the result of these reversions. We can see that:

**Table 1:** Observation and analysis of revertant colonies: A-Observation and phenotypic analysis of revertant colonies from mit- mutants M3041, M2075 and W91; **B-** Observation, phenotypic and molecular analysis of revertants of the CK247 mutant recorded in three experiments 1, 2 and 3.

|  | Type of reversion | Presence of sectors |
| --- | --- | --- |
| **CK247 (M3041)** | 75 % “Delta” | 15 % Sectoried |
| **CK312 (M2075)** | 100 % “Long” | 0 % Sectoried |
| **CK239 (W91)** | 100 % “Long” | 0 % Sectoried |

**B**
| Colony Phenotype | Number of CK247 revertants |  |  |  | Mol Genotype of revertants tested * |
| --- | --- | --- | --- | --- | --- |
|  | 1 | 2 | 3 | Total |  |
| **Large** | 119 | 75 | 65 | 262 | 82/84 Delta |
| **Large Sectoried** | 21 | 16 | 13 | 50 | 39/39 Delta |
| **Small** | 41 | 25 | 24 | 90 | 28/28 Long |
| **Small Sectoried** | 0 | 0 | 1 | 1 | 1 Delta |

\*: Among all the revertants of M3041, a certain number of revertants were analyzed by *in situ* hybridization using a probe carrying the bi2 intron and by HhaI RFLP analysis (see fig. S1).

- CK239 (W91) and CK312 (M2075) revert almost as frequently as CK247 (M3041) (that is to say 3 to 5.10^-8^) and show no sector on their revertant colonies. In total, more than 200 revertants per mutant were analyzed.

- All but one of the sectors appeared only on large colonies of CK247 revertants and all appear to be intron-deleted ("delta" type) and represent 15% of the large colonies. A number of M3041 revertants were analyzed either by *in situ* hybridization using a probe carrying the bi2 intron or byrestriction analysis of total DNA using HhaI enzyme (29), see fig. S1.

-The exception came from a small colony with a sector but which turned out to be of the delta type!, in short, *the exception that proves the rule*.

- The twenty-eight other small colonies, analyzed molecularly, proved to be revertants having retained the bi2 intron. Note that a small colony (< 2 mm) contains on average 10^6^ to 10^7^ cells.

These results strongly suggest that the sectors emerge exclusively after reversion by mitochondrial intron deletion. The relationship (intron deletion – sector) can therefore be suggested but not definitively established. Indeed, one can argue against this and say that the chance of seeing a sector on a small colony is lower than on a large one, despite the existence of the only small sectored colony. However, other arguments will later reinforce the idea of a probable relationship between the two phenomena.

### Physiological characterization of S and AS strains

The physiological analysis of the different cell types observed should make it possible to specify how the S cell is different from AS?. Do all S have the same physiological properties?. To try to answer these questions, two sectored colonies were chosen and analyzed. Colony 2 corresponds to a delta-bil+bi2+bi3 type revertant, and colony 3 corresponds to a delta-bil+bi2. The two parts AS and S of each colony were carefully picked, cultured separately, and designated AS2, S2, and AS3, S3 respectively. They were then cultured in two liquid-rich media: N3 (Glycerol) and YP10 (Glucose) and their growth kinetics were characterized. On the Glucose medium, there was no difference between the AS and S clones while a large difference was observed on the Glycerol medium (N3) between the AS and S clones, see Table 2. Indeed, the Optical density measurement made it possible to show that the growth of S cells was characterized by a shorter generation time than that of “AS” clones: 3h35 for S2 compared to 4h for AS2. This difference reflects the sector protrusion observed in Fig. 1A-C. S growth was also characterized by a lower value of the stationary phase, which reflects the subsidence of the sector compared to the rest of the colony (Fig. 1A-C). Indeed, the stationary phase of S is reached at an OD equal to half that of AS: 240 and 420 respectively, Table 2. The supplementary figure fig. S2 summarizes these differences and shows the correspondence between the growth curves and the shape of the sector and the rest of the colony. Indeed, the protrusion of the circumference results in the difference between the two growth phases (indicated by a red line) while the subsidence of the sector compared to the rest of the colony results in the difference between the two stationary phases (indicated by a blue line), Suppl Fig.1. Note that the growth curve of S2 is identical to that of S3 and that of AS2 to AS3.

**Table 2:** Physiological characteristics of AS and S strains. AS2 and S2 strains were grown in N3 (Glycerol) and YP10 (Glucose) and the kinetic parameters are presented in the table: generation time (GT) indicated in minutes and stationary phase (SP) shown in OD Klett measurements. The values presented are the average of three experiments. Note that strains AS3 and S3 give approximately the same results as AS2 and S2 (data not shown). The *p*-values indicated are according to Fisher’s test (0.0003; OR=0.6379) and Chi 2 test (0.9672; OR=0.9940).

| Strain | N3 |  | YP10 |  |
| --- | --- | --- | --- | --- |
|  | GT | SP | GT | SP |
| AS2 | 240 | 420 | 85 | 1335 |
| S2 | 215 | 240 | 85 | 1327 |
| *p* | **0.0003** |  | 0.9672 |  |

We therefore sought to see if several S and AS strains presented the same growth profiles, simply by measuring the OD of the stationary phase. Therefore, multiple S and AS cells from 16 different CK247 (M3041) revertant colonies were collected and cultured in N3 medium until stationary phase. The extreme limits of OD obtained by category of revertants (the values being the average of three different experiments) are as follows: AS = 24 - 32 and S = 16 - 22. We observe that the OD of the stationary phase is characteristic of each category, with a very significant difference between the S and AS strain pools, with *p* = 0.0001 (Fig.2). Growth on a rich medium containing glucose did not reveal any difference between the CK0/1, AS2, AS3 or S2, S3 strains either on the generation time or on the stationary phase. The physiological relationship of this phenomenon of *sectorization* with respiration/mitochondria is therefore suggested.

**Fig. 2:**
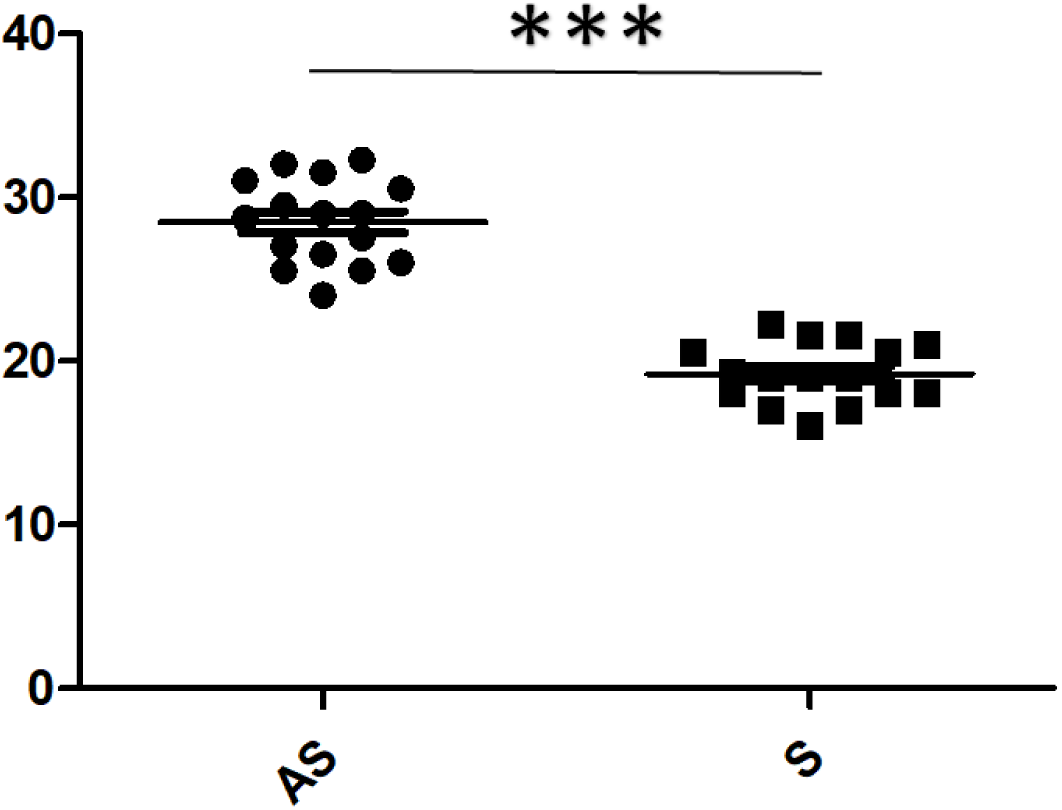
Optical density (OD Klett) in stationary phase of different CK247 revertants. 16 different revertant colonies phenotyped as sectored, from which AS and S cells were isolated. AS and S were then cultured in N3 liquid medium for 3 days at 28°C with shaking. The O.D Klett is measured after a 1/20 dilution of the culture. The values presented are the average of three different experiments. The *p* value is 0.0001 according to Mann Whitney non-parametric T-test.

### Genetic determinism

We come to one of the obvious questions to ask: are the physiological states S and AS hereditary? What is their genetic determinism?

The demonstration of the inheritance of states AS and S is already underlying in what precedes. Nevertheless, we can affirm it by emphasizing that the descendants isolated by sub- cloning of the parents S2, AS2, S3 and AS3 (previously cultured for more than twenty generations) always keep the same OD in the stationary growth phase in N3 medium. Similarly, the spreading of AS or S cells always gives, after 5 days of incubation on N3 and at 28°C, colonies which are all identical to one another. There is no heterogeneity between the colonies on an average of 200 colonies per strain tested.

After having suggested that sectorization is linked to the deletion of mitochondrial introns during the reversion of intron mutations, one is tempted to say that the cellular state S would be mitochondrial/cytoplasmic inherited (known as maternal in higher organisms).

To test this hypothesis, cytoduction is the method of choice. Indeed, taking advantage of the fact that these strains possessed the Karl-1 mutation, it was possible to carry out the cytoduction of mitochondria other than AS or S into rho^0^ cells containing AS or S nuclei (cytoduction A, Table 3A) as well as the cytoduction of AS or S mitochondria in another nuclear context (cytoduction B, Table 3B).

**Table 3.** : Optical density. (OD_Klett_) at the stationary phase of the different haploid strains constructed by Cytoduction (tables A and B). All strains constructed are incubated in liquid N3 medium for 3 days at 28°C with shaking. The OD_Klett_ is measured after a 1/20 dilution of the culture. The name of the mitochondria “donor” strain is indicated (Mitochondria) and the name of the "recipient" rho^0^ strain is indicated (Nucleus).The *p* values were calculated according to the unpaired T-test and according to Krustal-Walis Anova nonparametric test.

3A- CYTOFUSION A
| Mitochondria | Nucleus |  |  |  |
| --- | --- | --- | --- | --- |
|  | AS2 | S2 | AS3 | S3 |
| AS3 | 32.0 | **18.0** | 30.0 | **18.0** |
| W303-1B | 37.5 | **21.0** | 29.0 | **20.0** |
| 777-3A | 47.5 | **26.0** | 45.0 | **19.0** |
| *P* | 0.0273 |  | 0.0396 |  |

3B- CYTOFUSION B
| Nucleus \ Mitochondria |  |  |  |  | *p* |
| --- | --- | --- | --- | --- | --- |
|  | AS2 | S2 | AS3 | S3 |  |
| **W303-1B** | 29.0 | 32.0 | 30.5 | 30.8 | 0.3916 |

To carry out the cytoduction-A, the two pairs of strains AS2, AS3, and S2, S3 (all carrying the Kar1-1 mutation, required for cytoduction, as explained before) were converted into rho^0^ clones by mutagenesis with ethidium bromide. A rho^0^ clone of each strain was then sub-cloned and used for the following crosses, it will be called OAS or OS depending on whether it is a rho^0^ AS or a rho^0^ S. Several crosses were made between these rho^0^ and the following Gly+ strains: 777-3A, W303-1B and the previously constructed cytoductant W303-1B/AS3. Several cytoductants having the different mitochondria associated with the AS and S nuclei were isolated and sub-cloned and subjected to the liquid growth test in N3 medium.

To carry out the cytoduction-B, the rho^0^ W303-1B/50 strain was crossed with the various parents AS2, S2, AS3, S3 and the Gly+ cytoductants having the W303-1B nucleus were selected and sub-cloned.It should be remembered that both strains carry the Kar1-1 mutation, responsible for the delay in karyogamy and essential for the achievement of cytoduction. It should also be remembered that each parent carries a genotype that is easy to distinguish from the other: W303-1B (α) leu2, trp1, ura3, his3, CanR (31) while the parents with the sign (a) have the genotype (a) leu1, CanR, Kar1-1 (31).

Analysis of cytoduction results (Table 3): It should be noted that the test simply consisted in inoculating the different strains constructed in N3 medium at low cell density (10^4^ cells/ml) and incubating the shaken cultures for 3 days at 28°C. At this term, the stationary phase is reached and the OD at the end of growth is then measured.

This analysis reveals that it is the nucleus that determines the physiological state S or AS and this, is in the presence of any mitochondria rho^+^mit^+^. Indeed, Table 3A shows that the cytoductants possessing the S nucleus and having received various mitochondria, presented the lowest OD in the stationary phase than their AS counterparts, in a very statistically significant manner. This was validated twice since we analyzed the results relating to clones with the nucleus from AS2 and S2 (*p* = 0.0273)as well as AS3 and S3 (*p* = 0.0396). On the other hand, the cytoductants possessing the W303-1B nucleus and having received mitochondria from clones S2, S3, AS2 and AS3 showed no statistical difference in their OD value in stationary phase (*p*=0.3916), Table 3B.

It can therefore be said that there is a nucleus in S cells which is different from the nucleus in AS cells. In summary, we are therefore in the presence of a phenomenon of genetic instability, triggered by a deletion of mitochondrial introns, which generates a particular physiological state whose inheritance is nuclear.

### Another piece of the puzzle

We wondered if the state AS is a necessary and sufficient condition to generate the state S?. On nearly 300 AS cells plated at 30 cells/N3-plate and 50 cells plated at lower density (less than 5 cells/plate); after 10 days of incubation, no sector was visible on the colonies obtained. We should have at least 50 sectorized colonies, if the AS state was necessary and sufficient to generate the S state at all times. This result is both important and surprising since it implies a temporal relationship. Additionally, CK0/1, S2 and S3 strains were also propagated. On the same number of colonies (300), no sector was detected. This result is relatively understandable since the strain S is already of type S and CK0/1 would not possess the still unknown genetic element, of the AS nucleus, supposedly necessary to be able to generate the sector.

What then is the other new element necessary to convert the cell into an S status, an element which would only be present during the first generations following reversion by intron deletion?… If we again admit the relation of (intron deletion – sector), one can think that the mitochondrial intron, after its deletion, would pass into the nucleus to cause the state of the sector there. In order to test such hypothesis of the transfer of mitochondrial introns into the nucleus during the sectorization phenomenon, the total DNA of the S2 and S3 strains was extracted and purified on a ClCs gradient. The three bands corresponding to mitochondrial, nuclear and 3μ plasmid DNA were purified. The analysis of the nuclear DNAs and cut by the *Hhal* or *EcoRI* enzymes showed, after transfer on gel and hybridization with purely intronic probes (bil and bi2), no specific hybridization even after long exposure. The same result was confirmed with the total DNA extracted by the minilysatemethod (43) from the S2 and S3 strainsand hybridised to the same probes. There is no trace of bi2 intron in the S nuclear background.

### Macroscopic chromosomal rearrangement

The second molecular study of the sectorization phenomenon concerned the analysis of the electrophoretic karyotypes of the AS and S strains. Indeed, the appearance of the S state could result from a chromosomal rearrangement such as a translocation affecting, for example, one or more chromosomes with or without the intervention of the mitochondrial agent. To test this hypothesis, the analysis of the chromosomes of the strains AS2, AS3, S2, and S3 as well as those of the wild strains CK0/1 and mutant CK247 was carried out by the method of electrophoresis in agarose gel in pulsed fields.

The result of the PFGE (Pulsed-Fields Gel Electrophoresis) analysis allows us to draw a conclusion that is as interesting as it is unexpected (Fig. 3):

**Fig. 3:**
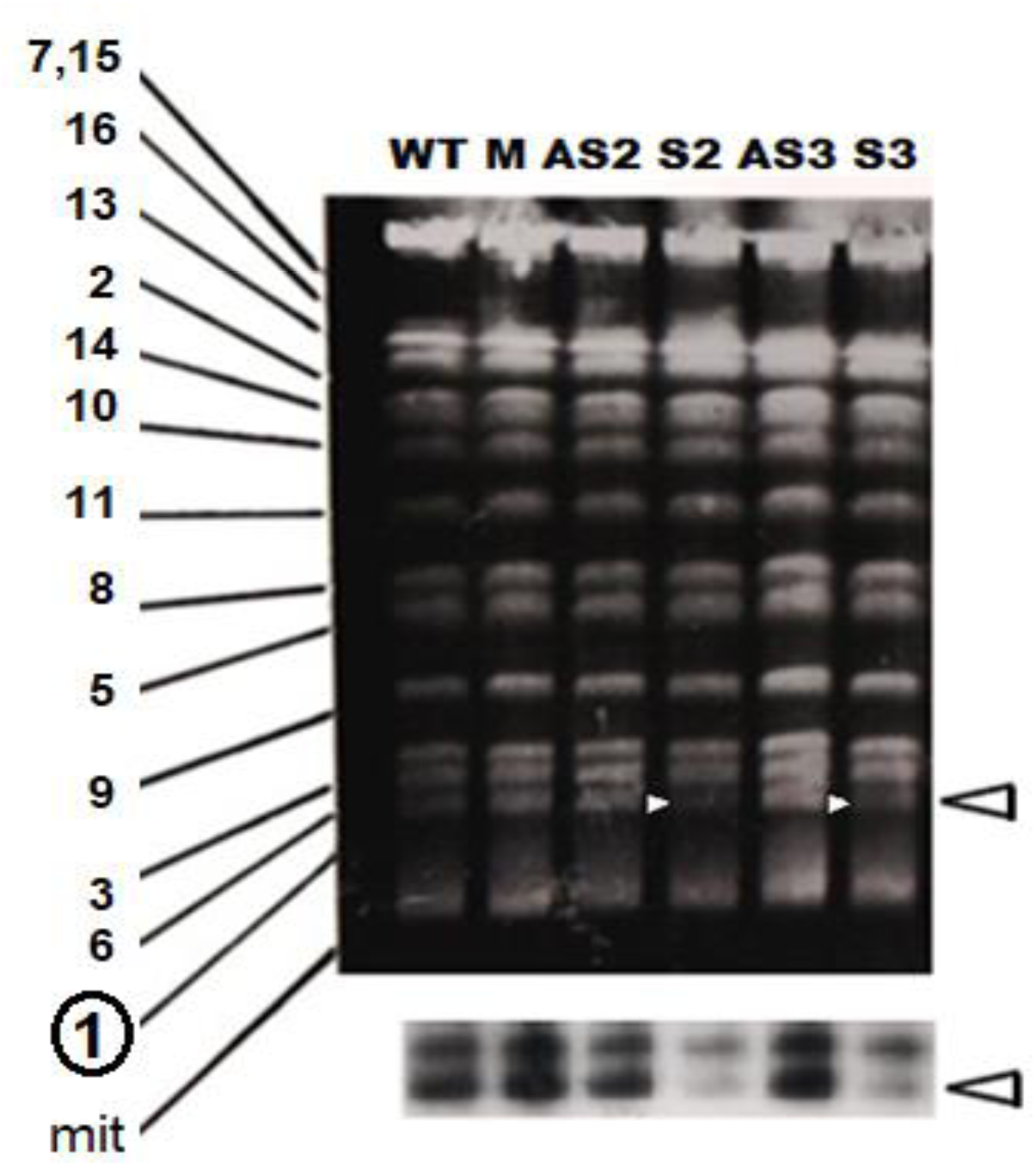
Pulsed-field gel electrophoresis analysis of chromosomes from S and AS strains and controls. Photo of the PFGE gel colored by BET, with WT: wild-type strain CK0/1, M: mutant strain CK247, AS2: anti-sector of colony 2, S2: sector of colony2, AS3: anti-sector of colony 3, S3: sector of colony 3. Left: chromosome number and mitochondrial DNA (mi). Bottom: The result of the hybridization of the transfer by the YCP:(ADE1) probe. The arrows indicate the position of chromosome 1.

The two S strains are distinguished from the others by the intensity of the band corresponding to chromosome 1 while in all the other control or AS strains, all the chromosomal bands have the same intensity, which is expected for a normal haploid or diploid cell with the same number of each chromosome. An aneuploid status with one chromosome less must present a less intense band than the others. This is precisely the case of the S2 and S3 strains where the band of chromosome 1 has a lower intensity than the other bands (Fig. 3). The densitometry of the gel bands shows that the ratio of the quantity of chromosome 1 ratio to that of another chromosome is close to 0.5 in the S2 and S3 strains and close to 1 in the AS2, AS3, CK247 or CK0l/1 strains. The hybridization of the PFGE transfer was carried out using a plasmid probe specific for chromosome1 (YCP:(ADE1)). The probe hybridized with chromosome 1 giving a much weaker signal in the clones of sectors S2 and S3 confirming their aneuploidy status, Fig.3. Note that cross-hybridization has been observed with some chromosomes, including chromosome 3.

Several hypotheses have been put forward to explain this molecular state is linked to the status physiological and genetic S.

1^st^ hypothesis: The S type cell would be diploid for 15 chromosomes and haploid for chromosome 1, ie: 2 x CH2-16 + 1 x CH1. It would therefore be monosomic aneuploid. However, the parent mutant strain CK247 (M3041) is normally haploid (a). It must therefore be assumed that during its lifetime, CK247 must have diploidized spontaneously and become (a/a). It cannot have become (a/α) since we have already seen that the AS and S cells could be crossed with a cell of sex sign α, cytoduction was even possible. In summary, the AS cell would be a true diploid (a/a) which, by losing a copy of chromosome 1, would generate an S sector made up of aneuploid cells.

2^nd^ hypothesis. The S strain would be a composite of normal haploid cells (a) and haploid cells (a) having lost chromosome 1. The latter would logically be non-viable since this chromosome, although it is the smallest: 260 Kb long, contains several genes essential for cellular life!. However, the extracted DNA would represent the resultant of two cell types, including *cadaver* cells without the CH1. As a result, the band of chromosome 1 would be less intense. If this hypothesis is true, the mortality would be twice as high in the case of the S strain. However, the mortality rate is not lower than that of the other control strains (data not shown). The hypothesis is therefore rejected.

3^rd^ hypothesis. The S strain is haploid (a) but heterogeneous with cells with all their chromosomal complement including chromosome 1 (in normal quantity) and cells with chromosome 1 in a different state. This state could be of two kinds: a) chromosome 1 would have circularized; b) chromosome 1 would have been entirely translocated to another chromosome. However, the absence on Profile S of a new band or change in the size of the other chromosomes does not support these hypotheses. Moreover, the subcloning of the S strains should have given clones different from each other, under one of these hypotheses. Also note that dicentric chromosomes are known to be unstable (32). It should also be noted that the hypothesis of a classic translocation (reciprocal or not) of part of chromosome 1 must also be ruled out since, in this case, the intensity of the band of chromosome 1 should remain unchanged.

Given these results, hypothesis 3, already unlikely, is ruled out while hypothesis (1) is consolidated; namely that the S strains are aneuploid by loss of a chromosome 1.

It was said above that -the result of the pulsed field analysis allows us to conclude as interesting as it is unexpected-. We said –unexpected- because our CK247 strain, in which sectors were detected, is normally a haploid strain whereas we now suspect it to be a diploid strain and aneuploid for chromosome 1!. This strain would have accidentally become diploid (a/a) during subculturing and previous cultures.

### Genetic test of the hypothesis

The analysis of the products of meiosis resulting from the crossing of the S and AS strains with an alpha haploid, marked differently at the level of chromosome 1 and of the other chromosomes, should make it possible to verify the hypothesis of aneuploidy as described in the preceding paragraph.

It should be recalled that the S and AS strains are derived from a haploid strain of genotype (α) leul can^R^ Karl-1. The recessive markers leul and can^R^ are respectively on chromosomes 7 and 5. Since chromosome 1 carries the Adel gene, an easy-to-interpret cross would be with strain 777-3A carrying precisely the recessive adel marker. In addition, this strain carries another recessive marker gene op1 located on chromosome 2. The AS and S strains, fortunately, carry the wild-type Adel+ and op+ alleles.

The prediction would be to obtain, based on the aneuploidy hypothesis and as it is described and explained in Fig.4 the following situation during the mass analysis of the spores of the crosses S x 777-3A and AS x 777-3A. Let us note before commenting on the results of this experiment that the method of analysis of the spores in mass was retained rather than that of the tetrads for the following reason. At the very beginning of this work on "sector colonies", thirteen tetrads were analyzed from the AS2 x W303-1B/50 cross and twelve tetrads from the S2 x W303-1B cross. In both cases, more than 75% of the spores were not viable. This initially misinterpreted result becomes logical since it is well-known and plausible to have very poor spore viability in the case of meiotic segregation of a triploid (33). It would therefore have been necessary to analyze a gigantic number of tetrads from the 777-3A x AS2 and 777-3A x S2 crosses to have a suitable number of spores for the statistical analysis. Mass spore analysis, on the other hand, makes it possible to have a large number of spores without any difficulty. It is however mandatory, in this kind of analysis, to have genetic markers to distinguish a colony resulting from a diploid cell from a colony resulting from a spore. This distinction is in our case possible and very easy thanks to the markers involved and in particular the recessive marker Can^R^. Indeed, in addition to the W0- colonies (auxotrophic for at least one of the ade1 or leu1 markers), all the W0+ Can^R^ colonies (W0+ means prototroph) are automatically considered as resulting from spores. Only (W0+ Can^S^ Gly+) colonies must therefore be eliminated at the risk of confusing them with a colony resulting from a diploid.

**Fig. 4:**
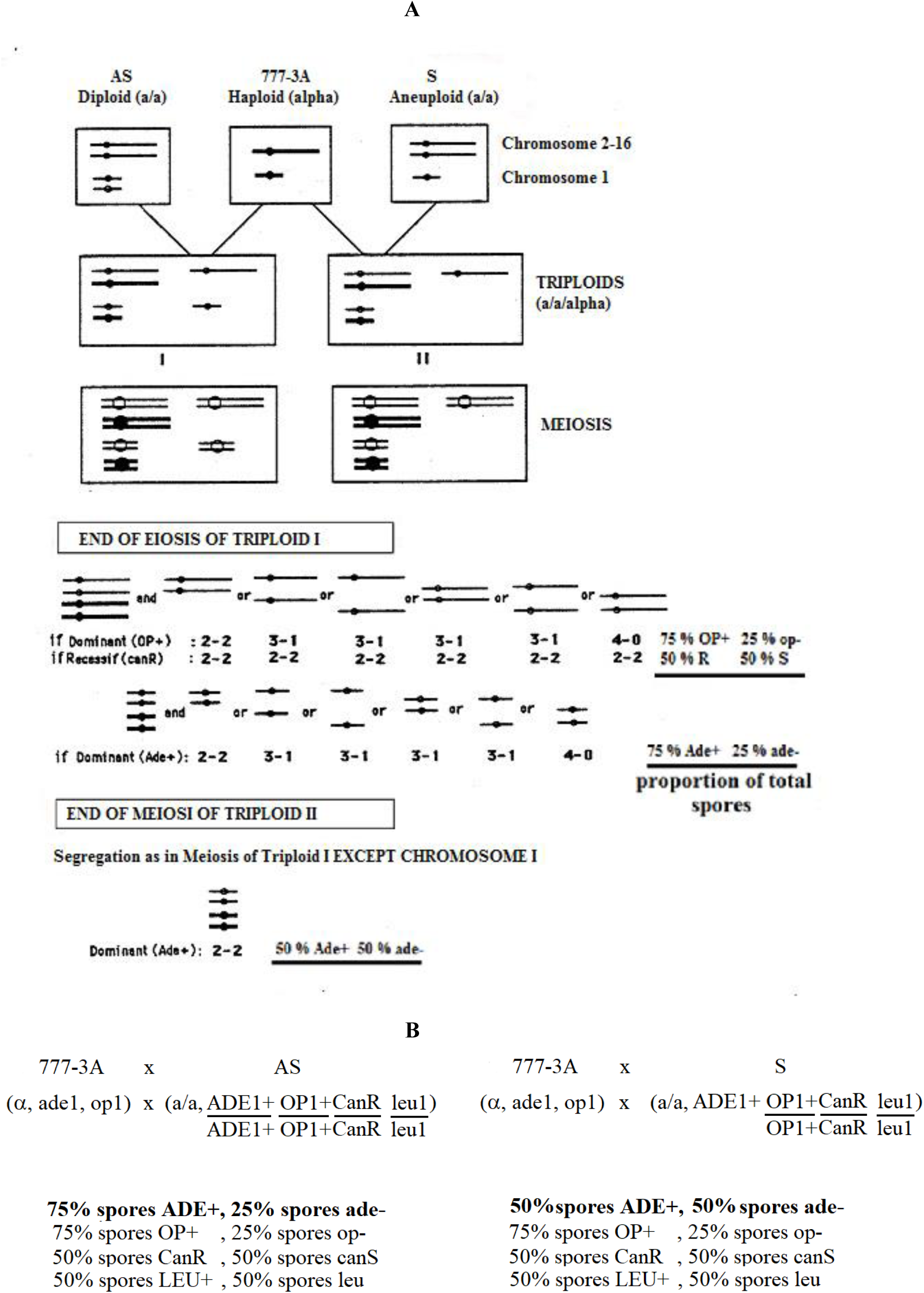
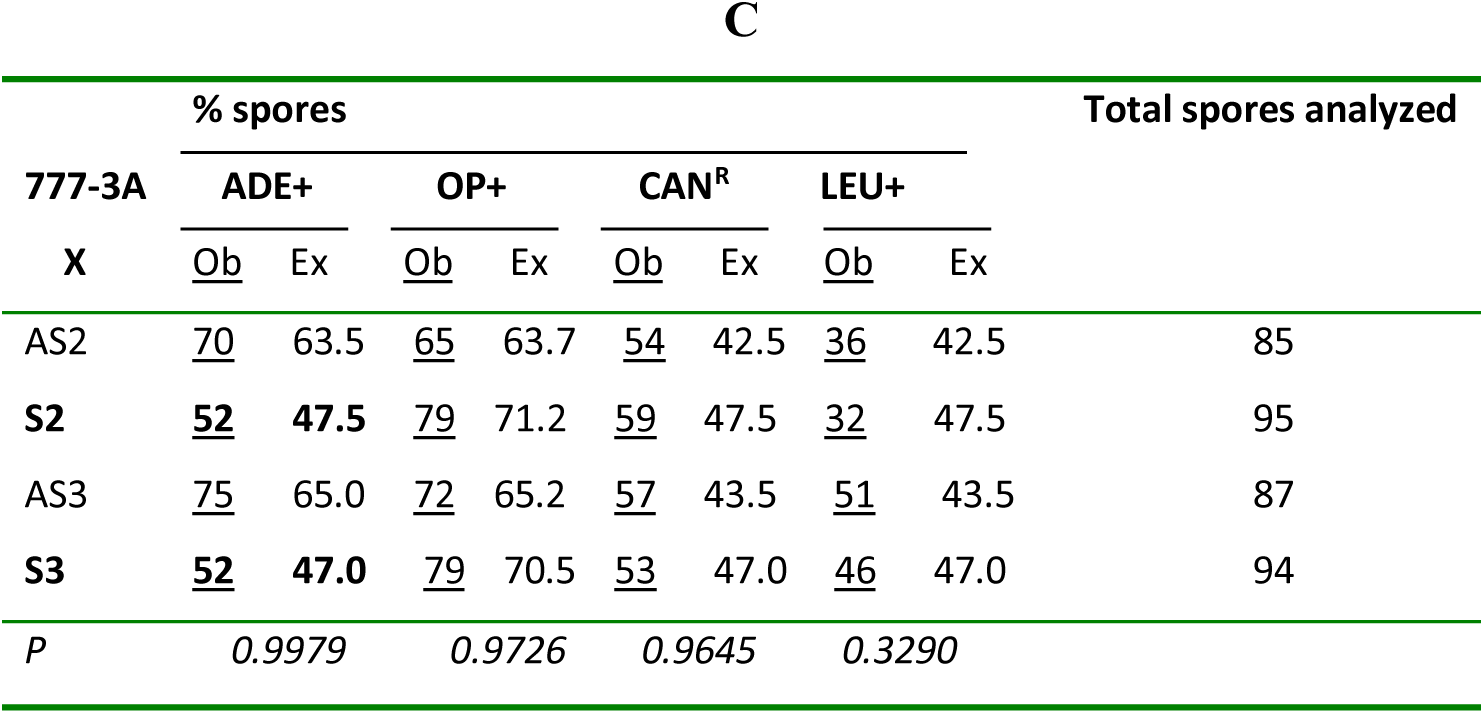
Schematic of the genetic analysis of meiotic segregation of triploids resulting from crosses AS x 777-3A and S x 777-3A. **A-**For each cell, a single representative of chromosomes 2 to 16 is represented by long horizontal segment. Chromosome 1 is represented by the short segment. The different possible combinations of supernumerary chromosomes in tetrads are presented at the end of meiosis of triploid I (AS x 777-3A). These combinations are also valid for meiosis of triploid II (S x 777-3A) with the exception of chromosome 1. **B-** The genotypes of the cross strains are shown at the top, the expected proportions of spores resulting from triploid sporulation, according to the analysis shown in Figure 3A, are shown below. **C-** Are indicated in the table: Ex: The expected number of spores per phenotypic class, Ex, is calculated based on the aneuploidy assumption (see Fig. 3A) and the total number of spores analyzed per cross; Ob: The number of spores actually observed is indicated by Ob and underlined. Note that the markers Ade, Op, Can and Leu are specific of chromosome 1, 2, 5 and 7 respectively. The *p*-values are shown in the last row of the table, according to the χ2 analysis.

The table in figure 4 (Fig. 4C) shows clearly that the proportion of ade+ spores resulting from meiosis of the S2 x 777-3A or S3 x 777-3A cross becomes equal to 50% whereas it was about 75% in the AS2 x 777-3A or AS3 x 777-3A cross, as predicted based on our hypothesis modeled in Fig.4A. The proportion remains almost the same for Leu+, op+ or Can^R^ in both crosses, as also expected. The table indicates that the observed result is in perfect agreement with the theoretical prediction formulated above, according to the statistical analysis, which shows that the difference between Ex (Expected number of spores) and Ob (Observed number) is not significant.

This genetic analysis confirms therefore the prediction based on the molecular analysis: AS is an (a/a) diploid and S is also a/a diploid but aneuploidy for CH1, so having 2 copies of CH2-16 and one single copy of CH1.

### The *return question*: Is the sector genetic background favorable to reversion by deletion of mitochondrial introns?

The question posed is the following: if reversion by deletion of mitochondrial introns favors the appearance of sectors, is the nuclear background of the sector, in turn, favorable for the deletion of mitochondrial introns?

The approach followed, shown schematically in Fig.5, is summarized here:

**Fig. 5:**
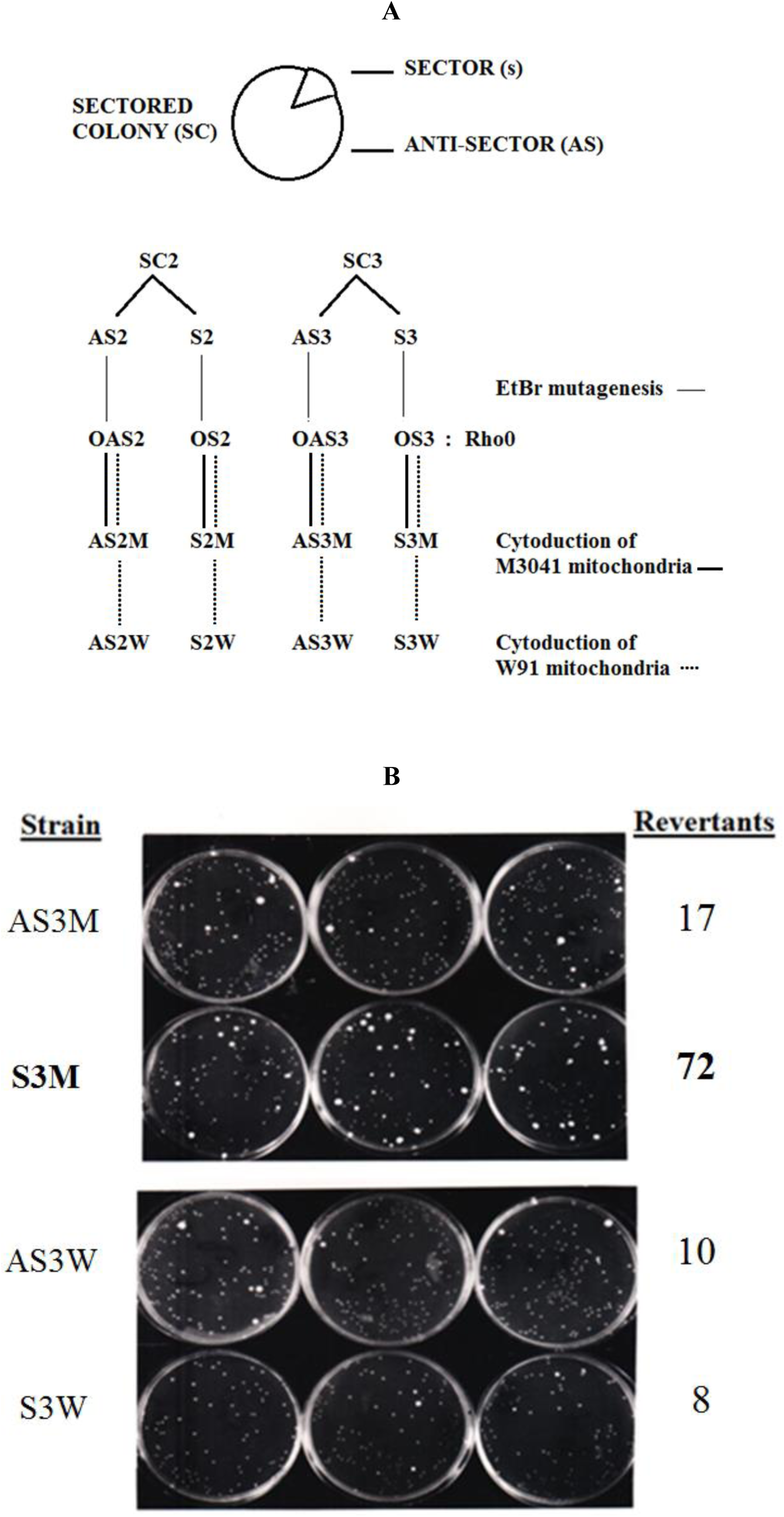
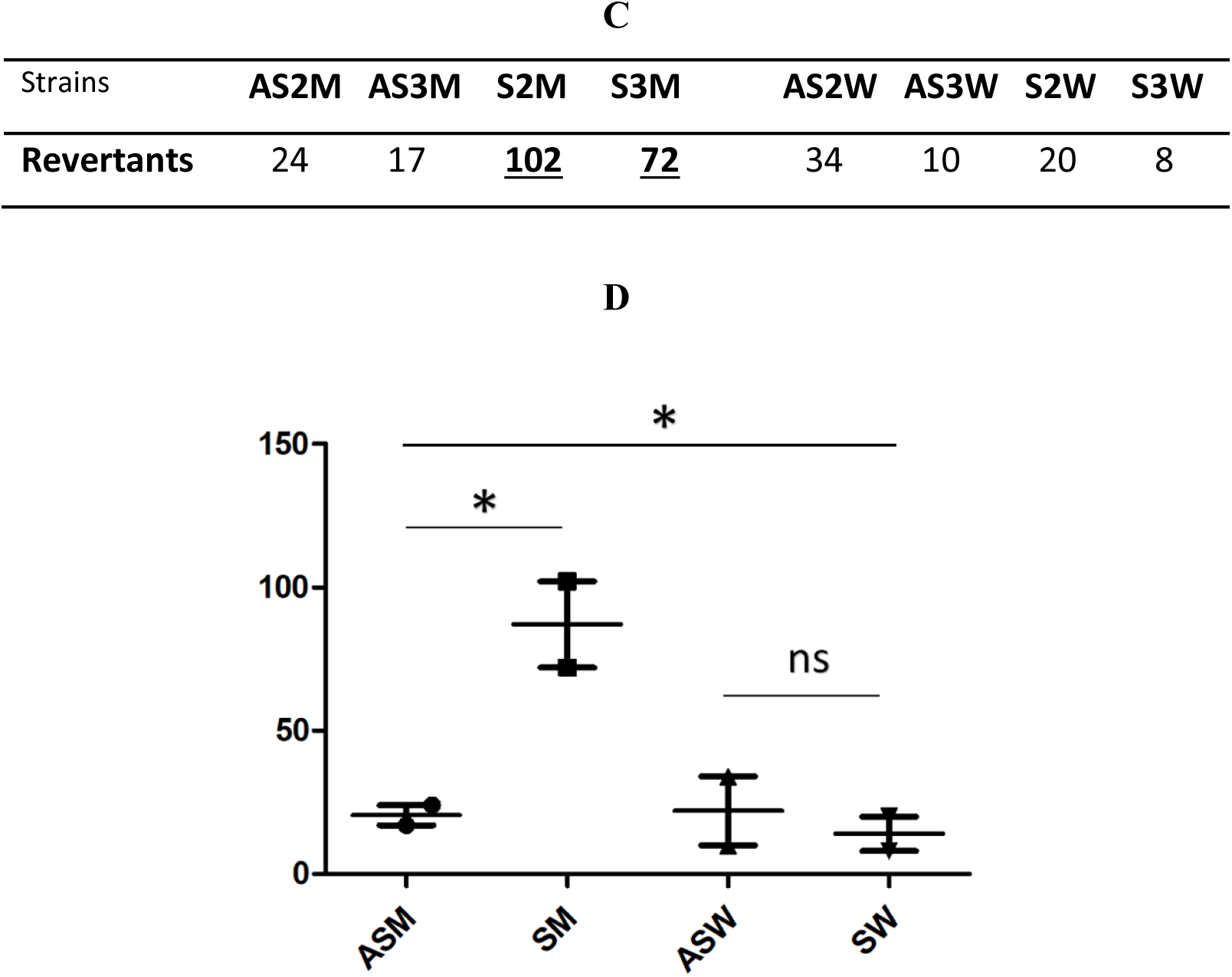
Return relationship -sector to delta-: **A.** Construction strategy: two sectored colonies of CK247 revertants (a/a, leu-) were chosen. They were different as SC2 had a deletion of bi1+bi2+bi3 whereas SC3 had a deletion of bi1+ bi2 introns. The anti-sectors AS and sectored cells S were picked and converted to rho^0^ by EtBr mutagenesis and named OAS2, OAS3, OS2 and OS3. After verification of their rho^0^ status, they were crossed to M3041 or W91 strains (α, ade-). The cytoductants of (a/a) nuclear backgrounds were selected and checked for their mitochondrial background and named accordingly: AS3M = M3041 cytoducted in the rho^0^ OAS3; S3M = M3041 in the rho^0^ OS3; AS3W = W91 in the rho^0^ OAS3; S3W = W91 in the rho^0^ OS3. Then, they were allowed to revert on YPDif plates, an example result of which is shown in the following picture. **B.** Photographs of the YPDif plates of the revertants of the AS3M, S3M and AS3W, S3W strains (knowing that clones AS2M, S2M and AS2W, S2W gave a similar result). The names of the strains plated are shown on the left. On the right: the numbers of revertants recorded. **C-**The values represent the number of independent revertants obtained after incubation of the cells of the various strains on YPDif for 15 days at 28°: AS2M, S2M, AS3M, S3M relative to M3041 as well as AS2W, S2W, AS3W, S3W for W91. **D-**Graph of the table presented in C, The *p* values were: 0.0203 according to One-way Anova for all values; 0.0497 for AS2M, S2M, AS3M, S3M values and 0.6115 for AS2W, S2W, AS3W, S3W, according to Unpaired-test.

- Cytoduction of cytoplasm/mitochondria of M3041 in rho^0^ OS2, OS3, OAS2, and OAS3 as test mutation of "intron deletion reversion" and mitochondria W91 as negative control since W91 never reverts by intron deletion (28,29).

- Reversion test on the strains constructed by spreading a hundred cells per YPDif dish to estimate the frequency of reversion by the method already described (29).An example of the plates is given in Fig.5B with the result summarized in the table below (Fig.5C).

We counted the apparent revertants on almost the same total number of "mutant" colonies formed on YPDif medium (between 330 and 390 on the total plates, by construction). There were approximately four times more independent revertants in the case of the S2M and S3M strains (M3041 associated with the S2 and S3 nuclear contexts respectively) than in the AS2M and AS3M strains (M3041 in the AS2 and AS3 contexts), (Fig. 5C), the difference being statistically significant with *p*= 0.0497. In the case of the control strains (having W91), the frequency of reversion is substantially the same in the four constructions S2W or S3W and AS2W or AS3W, with *p*= 0.6115 (Fig. 5C). Fig.5B shows the example of results obtained with the AS3 and S3 backgrounds. Note that the reversion frequency per cell for the AS2M, AS3M, AS2W, S2W, and S3W strains is close to 3.l0^-8^, a frequency comparable to the diploid strains KM716 (M3041) and KM109 (W91) as well as the haploid strains CK247 (M3041) and CK239 (W91) (27–29).

20 revertants from each construct were analyzed by *in situ* hybridization using the probe containing the bi2 intron and confirmed, for some of them, by restriction analysis using *HhaI* enzyme as shown in fig. S1 and in (29). We recall that this simple technique allows us to distinguish long and delta revertants by *HhaI* restriction analysis (see fig.S1) from their total DNA extracted by a simple mini-lysate procedure (27–29). It is preferred to PCR analysis for its simplicity and for the fact that the deletion of different introns should lead to different lengths of PCR fragments (of several kb) depending on the number of introns deleted. All revertants from constructs AS2M, and S2M (containing M3041) were of "delta" type (with intron deletion) whereas all revertants from constructs AS2W, and S2W, were of “long” type (had not lost introns).

It can therefore be concluded that the aneuploid S nuclear background favors the deletion of the intron (or facilitates its detection) and has no effect on the other modes of reversion of M3041 mutation. On the other hand, the nuclear diploid AS background does not favor intron deletion. We can think that the dose of chromosome 1 plays an active role in the process of intron deletion and suppose that it carries a gene(s) which, when it is in a single dose, in front of other genes in a double dose, would cause some resultant nuclear expression to stimulate the deletion of mitochondrial introns.

## Discussion

This study shows an intriguing relationship between the reversion of mitochondrial mutations by intron deletion and nuclear aneuploidy; What’s more intriguing is that this relationship exists both ways. In the first part of this work, we reported the fortuitous discovery of a -sector on colony- phenotype in revertants of the M3041 mutant, only those who reversed by intron deletion. By analyzing two independent reverting colonies, by pulsed fields electrophoresis, we showed that the sectors corresponded in fact to a loss of the chromosome in an (a/a) diploid. If we put together the different pieces of the puzzle at our disposal, the following pattern seems to emerge:

- CK247 (M3041) would have diploidized by becoming (a/a)! AS would therefore be an a/a diploid which generated the sector by loss of chromosome 1.

- Sector occurs only after the deletion of mitochondrial introns.

- Sectors occur when one copy of chromosome 1 is lost. The cell then becomes aneuploid. It therefore has particular physiological properties, on glycerol medium (where only respiration could take place, fermentation being excluded) probably due to an effect of the dosage of the genes on CH1. S cells are thus physiologically deregulated.

- The S state is relatively stable, and is transmitted to descendants without apparent change.

- Having become aware of the genetic determinism of the S state, we understand why the CK239 (W91) and CK312 (M2075) strains could not give rise to a sector: they had not diploidized and had remained haploid (a).

However, several relevant observations remain:

1. AS cells no longer generate a sector when plated on N3 medium; Why?; It is as if the appearance of the sector was linked in time to the reversion by intron deletion.
2. CK247 long-revertants, i.e not having lost introns, were not able to yield a sector.

Two conditions would therefore have to be met for the appearance of the S state: (1) the strain must be a/a diploid (we do not yet know whether α/a diploids can give rise to sectors or not?); (2) Reversion by deletion of mitochondrial introns would cause or facilitate the loss of chromosome 1, by an unknown mechanism limited in time.

We can also speculate on this need to lose chromosome 1 as follows:

The deletion of mitochondrial introns (bi1 and/or bi2) generates a new mitochondrial genotype incompatible with nuclear-derived products. One can imagine that the absence of an intron or a maturase, for example, would make the presence of a specific product (encoded by a nuclear gene x) normally involved in RNA splicing harmful to mitochondria. Mutants no longer possessing this product would then be selected. This could be done in two ways: modification of this nuclear gene or elimination of a chromosome (here chromosome 1); the consequence of this elimination is to deactivate the gene x. Of course, for a haploid cell, the loss of a chromosome is fatal. In a diploid cell, it is possible and the genetic imbalance caused by this aneuploidization will be such that the mentioned incompatibility will no longer have any reason to persist.

If these speculations have any truth, one of their implications is that delta revertants would be more easily selected in an S-context.

This hypothesis was then tested in the second part of this work, in the so-called return- experiment. Indeed, we have shown that if an S and AS cell were converted into rho^0^ in which mitochondria M3041 or W91 were introduced by cytoduction. The number of revertants was significantly higher in M3041 in the S context than in AS, while it was the same for W91, the control strain who never reverts by intron deletion (28, 29). The M3041 revertants were mainly of the intron deletion or Delta type, which was absent in the W91 control. This means that the S context (aneuploidy for CH1) favors the deletion of mitochondrial introns during the reversion of mit- mutations.

In light of the results of the second part, we hypothesize that the loss of a single gene from chromosome 1 would be analogous to the loss of the entire Ch1. The mutation of this gene x by KO would be equivalent to the S genetic ground and becomes favorable for the deletion of the mitochondrial introns. To find this gene, we propose to implement the following strategy: from the bank of knockout yeast strains covering more than 6,000 individual genes, we select the mini-bank of Chromosome1 mutants, which contains approximately 100 KO mutants for each of these genes. This collection will first be converted into rho^0^ by EtBr treatment, then crossed with M3041strain. We will thus have a collection of diploids where each clone carries a mutated Ch1 gene in the heterozygous state, mimicking the S state (except that it is by inactivation of a single gene and not by loss of all the Ch1) and a mitochondrion carrying the M3041 mutation. The reversion test will then make it possible to identify the clone giving more independent revertants than the others. This contains the searched gene in a heterozygous mutated state. Of course, this approach can only succeed if -1- we accept that an (α/a) diploid can also give sectors like our starting strain CK247 which was (a/a), -2- heterozygosity can replace the aneuploidy, in other words, is the loss of a single Ch1 gene sufficient to obtain S status?

The sectors described here appeared in a strain which would have diploidized by becoming a/a. This is a kind of polyploidy that is actually quite common in vertebrates as well as in fungi and yeasts like *S. cerevisiae*. Indeed, beer strains are known to be polyploid, with two or usually more than two copies of their genome per cell (34). Polyploidy is also known to be associated with genomic instability, but the underlying mechanisms are not yet well understood (35). Polyploidy could be considered or explained as a solution allowing the yeast cell to escape the deleterious mutations that accumulate during replication in a haploid cell. According to Tracy et al. 2020, these mutations could be the consequence of defects in the exonuclease domains of DNA polymerase or mismatch repair (MMR) (36). These defects have been reported to cause tumor genesis in animals and humans. Cancers are the best example of this, with a large mutational load in addition to chromosomal rearrangements leading to polyploidy as well as aneuploidy and many other events (35).

Human oocyte aneuploidy has often been associated with advanced age in women, knowing that in men, this rate is ten times lower (37). Sonowal et al. 2023 showed that indoles prevent aneuploidy and promote DNA repair and embryo viability, which depends on age and genotoxic stress levels and affects embryo quality across generations (38). Although aging is closely linked to increased aneuploidy in oocytes, the mechanism by which aging affects aneuploidy remains largely unknown (38). Interestingly, some studies emphasize the relationship between aneuploidy and mitochondria, especially mitochondrial dysfunction. Indeed, Zhang et al. 2023 very recently showed that mitochondrial dysfunction and abnormal spindle assembly of aging oocytes can lead to increased oocyte aneuploidy (39). Another study showed, in the Drosophila epithelial model, that the aneuploidy-induced senescence may be the consequence of proteostasis failure and mitochondrial dysfunction (40). Even if this relationship between aneuploidy and mitochondria is very different from that revealed by our study, where there is no mitochondrial dysfunction, the fact remains that there is indeed a relationship between nuclear aneuploidy and mitochondria. In our case, the relationship between mitochondria and nuclear aneuploidy is not due to mitochondrial dysfunction, quite the contrary, it is a reactivation of mitochondria thanks to the deletion of introns.

In the first part, where the deletion of mitochondrial introns would lead to an elimination of chr 1, this elimination is related in time to the deletion of introns. What would be this molecule that would transmit this information from the mitochondria to the nucleus and cause the elimination of Ch1?.

Conversely, the second part of our study shows that the aneuploid genetic background would promote the loss of mitochondrial introns. Could this be a kind of activation of the reverse transcriptase activity necessary during the deletion of mitochondrial introns?. Remember that this activity is encoded by the two introns ai1 and ai2 of the mitochondrial cox1 gene and was shown to be homologous to viral reverse transcriptases (10, 11). Let us also recall that ai1 alone may be sufficient to achieve the reversion by intron deletion but that the presence of introns ai1 and ai2 is much more effective (29).

## Materials and methods

### Strains

All the strains used in this work derive mitochondrially from the strain 777-3A having the genotype: (a) adel, op1 (*41*). Origin of the CK247 strain: it has the mitochondria of 777-3A/M3041 and the JC8/55 nucleus. (JC8/55 is a rho^0^ (a) leu1, CanR, Kar1-1 (*31*)). M3041 is a trans-recessive mutation localized in the bi2 intron of the cyt.b gene (*8*). In order to eliminate the effect of op1 mutation (in the gene AAC2 (*42*) of the 777-3A nucleus), M3041 was passed by cytoduction in JC8/55 background and named CK247. Kar1-1 is a dominant mutation responsible for delayed karyogamy, useful in cytoduction; can^R^ confers the canavanine resistance phenotype.

CK247 is therefore a haploid strain with genotype (a, leu1, canR, Kar1-1) and carries the mit-M3041 in its mitochondrial DNA. CK239 and CK312 strains carried the 777-3A/W91 and 777-3A/M2075 mit- respectively, while CK0/1 carries the wild-type 777-3A mitochondrial DNA. Other strains used in this work: W303-1B which is a rho+, (α) leu2, trp1, ura3, his3, CanR (*31*).

### Media

YPG: 1% Yeast extract, 1% Bacto-peptone, 2% glucose. YP10: 1% Yeast extract, 1% Bacto-peptone, 10% glucose. N3: 1% Yeast extract, 1% Bacto-peptone, 2% glycerol. YPDif: 1% Yeast extract, 1% Bacto-peptone, 2% glycerol, 0.1% glucose (*22, 29*). Minimal Medium (W0L): 0.67% yeast nitrogen base, 2% glucose, to which were added 20 mg Leucine (L).

### Reversion on N3 plates

mit- mutants deficient in respiration, in the CK background (having the nuclear genotype of JC8/55), are cultured in YP10 medium without shaking until the stationary phase. Then, approximately 10^8^ cells are plated on each N3 or Glycerol plate and incubated for at least one week at 28°C. Few revertant cells emerge and are able to respire and consume glycerol until they form great colonies. Such revertants are then sub-cloned on N3 plates and analyzed by rapid DNA minilysate extraction (*43*) and analysis of their mitochondrial cyt.b gene.

### Independent revertants isolation on YPDif plates

This method ensures the selection of independent revertants allowing establishing the true reversion frequency of mitochondrial mutants (*29*). About hundred cells of a fresh culture of the mit- mutant were spread on YPDif plates and incubated at 28°C. On such medium containing 0.1% glucose and 2% glycerol, the respiratory-deficient cells formed small colonies by consuming first the glucose. These colonies stopped to grow after few days and are made up of 10^6^ to 10^7^ cells per colony. In some of these colonies, revertant papillae emerge and gradually grow by consuming the glycerol, until they reach a size much larger than the parent colony. These revertants are necessarily considered *independent revertants* since each comes from a different colony. These revertants are then sampled and sub-cloned on N3 plates and analyzed by *in-situ* hybridization (*29, 44*) or by rapid DNA mini-lysate extraction and HhaI restriction analysis (*29, 43, 44*).

### Ethidium bromide treatment

In order to eliminate the mitochondrial DNA leading to the formation of the rho^0^ strain, cells in logarithmic growth (at 10^7^ cells/ml) are treated with three cycles of culture in medium containing ethidium bromide EthBr (*45*). Such prolonged incubation ensures complete loss of mitDNA. The rho^0^ status is genetically verified by crossing the clones obtained with strains carrying mit^-^ mutations M3041, V277, V25, V276 taking place respectively in the cyt.b, cox1, cox2 and cox3 genes.

### Cytoduction

It is a mating-based technique taking advantage of the kar1-1 mutation that delays nuclear fusion (karyogamy).The strain whose mitochondrial DNA is to be changed is firstly converted to rho^0^ by ETBr mutagenesis (*45*) (see above) and then crossed with a strain of opposite sexual sign and which carries the target mitochondrial DNA. One of the two strains must carry the dominant Kar1-1 mutation. Remember that the Kar1-1 mutation delays karyogamy for up to nearly 8 hours; during this period, the first buds will emerge with one or other of the cytoplasm, hence the transfer of cytoplasm or “cytoduction”. In the hours following the crossbreeding by confrontation of the two strains (on solid or liquid YPG medium), different dilutions are spread on YPG plates and incubated for 3 days. The colonies obtained are characterized at the nuclear level, by replica on different selective media: MM to identify diploids, MMLC: minimum medium supplemented with leucine and canavanine (genotype of JC8/55) and finally MMA (minimal medium supplemented with adenine (genotype of 777-3A). The mitochondrial genotype is determined by crossing with rho- covering part of the mitochondrial gene of cyt.b such as rho- B111 and 231with B111 covering all of the second bi2 intron while B231 covering part of it (*8*).

### Digestion, electrophoresis, transfer and hybridization of DNA

Total DNA was extracted by minilysate procedure (*43*). Pure mit.DNA was extracted using ultracentrifugation on ClCs gradient (*27*). The digestion by restriction enzyme is carried out in the buffer recommended by the supplier but for the mini-lysate DNA, the restriction reaction buffer contained higher NaCl concentration (*27, 29*). Digestion of total DNA from the minilysate by HhaI allowed us to distinguish between long and delta revertants, see fig. S1. Indeed, the site recognized by HhaI, GCGC, is very rare in yeast mitochondrial DNA due to its low GC content (around 20%) whereas nuclear DNA is richer. Therefore the mitochondrial bands (and mainly that containing the cyt.b gene) are easily discernible at the top of the gel while the nuclear genes migrate as smears at the bottom of the gel (*27–29*), fig. S1.

Gel electrophoresis was carried out with 0.7% agarose in TAE (Tris–acetate–EDTA). 20 x SSC buffer was used during transfers of gels on nitrocellulose (44). Filters were hybridized overnight at 65°C in 6 x SSC, 0.1% SDS, 1 x Denhardt, and 100 ug/ml of sonicated Salmon-sperm DNA. Filters were washed at 65°C for 20 min in 2 x SSC, 0.1 SDS, three times and once in 0.1 x SSC.

Pure bi2 intronic probe was used: pYJL5, donated by C. Jacq, consists of the rho- B231 (fully included in intron bi2, (*8*)) cloned into the vector pBR322. Two probes specific for ai1 and ai2 introns were given by P. Netter. The radioactive labeling of the probes was carried out by nick translation (*44*).

### Pulsed-Fields Gel Electrophoresis

The treatment of yeast cells for pulsed-field electrophoresis was carried out according to the method developed by François Caron (personal communication): Yeast cells are mixed with agarose and injected as droplets in mineral oil. Thus, the agarose forms beads, encapsulating yeast cells between its meshes. The beads are then incubated in a PBS pH7 containing the zymolyase, for 1 hour at 37°C, to produce protoplasts. Subsequently, the protoplasts are lysed by adding EDTA and proteinase K. Under the microscope, one can follow all the steps and check that at the end, all the cells gently burst, ensuring that the chromosomes are undamaged. The beads are placed in agarose gel wells and run in a pulsed-field electrophoresis apparatus (*42*). Hybridization of the PFGE gel blot was carried out using specific chromosome 1 probe, YCP:(ADE1), kindly gifted by David Kaback (*47*), that contains the ADE1 gene (chromosome 1) and the markers TRP1 and CEN4 (chromosome 4).

### Statistical analysis

All statistical analyses were performed with GraphPad Prism software, using the Mann-Whitney U-test, One-way Anova, Unpaired-test as well as Fisher’s test and χ2-test, etc… Differences were considered statistically significant at p<0.05 (*), p<0.01 (**) and < 0.001 (***).

## Supporting information

spplemental figures

## Aknowledgements

This work is dedicated to the memory of Professor Piotr P. Slonimski who allowed me to carry out this work in his laboratory, as well as to the memory of Annie Lamouroux and Agnès Delahodde. I warmly thank Claude Jacq, Pierre Netter and David Kaback for the gift of the probes and Eric Petrochilo for the always interesting discussions. Many thanks to Nabil Zouari and Nizar Jammoussi for their help in formatting and editing the text; as well as to Raja Gargouri for her help in statistical analyses.

