## Supplementary material for "Intriguing relationship between chromosome I nuclear aneuploidy and mitochondrial intron deletion in yeast": spplemental figures

**Supplemental figures**

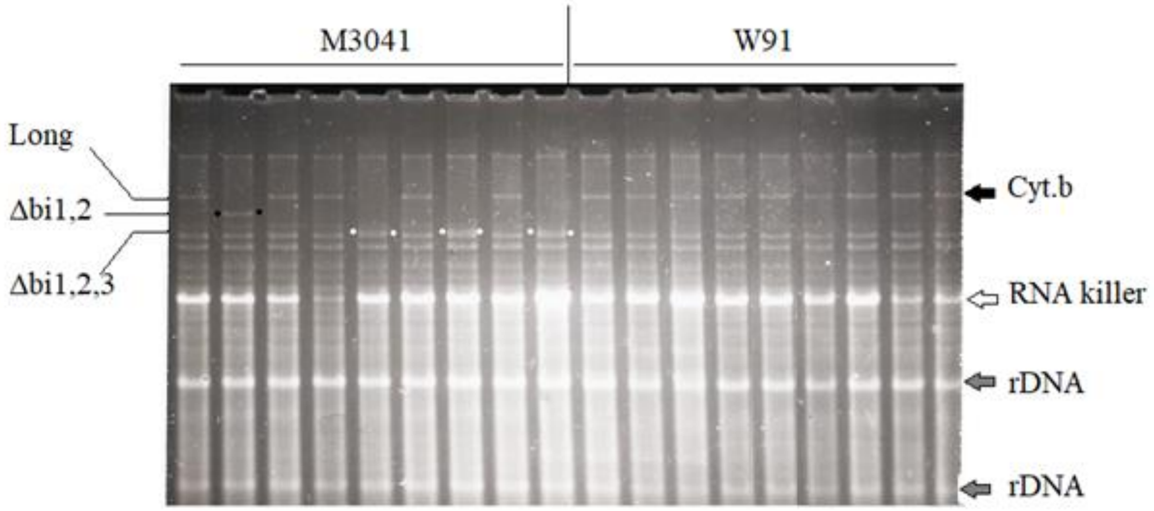


**fig. S1: Electrophoretic analysis of the DNA of the revertants by digestion with the restriction enzyme HhaI**. The name of the strains whose revertantsare analyzed is indicated at the top: M3041 and W91. On the right, the positions of the following bands are indicated: the fragment containing the entire mitochondrial cyt.b gene, the RNA killer, the nuclear rDNA fragments. On the left, the different sizes of the fragment containing the cyt.b gene are indicated: the gene with all the introns (Long Form), the gene deleted of introns bi1 and 2 (Δbi1,2, indicated by black spots); the gene deleted from introns bi1, 2 and 3 ((Δbi1,2,3 indicated by white spots). Note that the mit.DNA fragments are visible at the top of the RNAkiller while the nuclear DNA fragments form a smear at the bottom of the RNA killer.

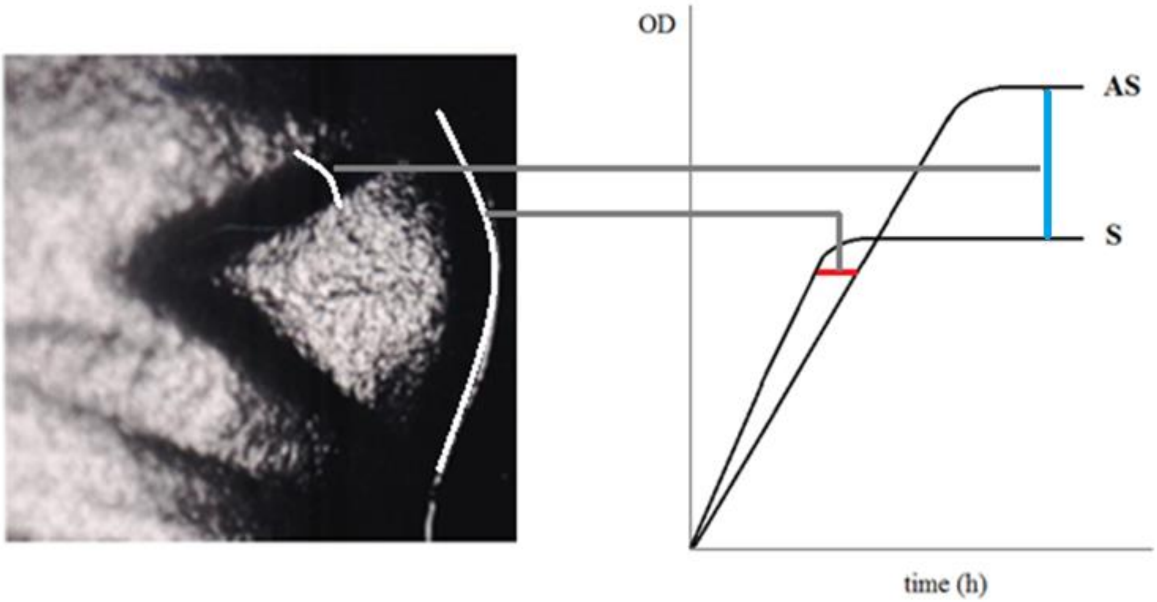


**fig. S2 : Correspondence between the shape of the sector and the growth curve of S (cells of the sector) and AS (cells of the rest of the colony).** The protrusion of the circumference is reflected by the difference between the two exponential phases (indicated by a red line). The subsidence of the sector in relation to the rest of the colony is reflected by the difference between the two stationary phases (indicated by a blue line).
